# Advanced Glycation End-Products Suppress Mitochondrial Function and Proliferative Capacity of Achilles Tendon-Derived Fibroblasts

**DOI:** 10.1101/706754

**Authors:** Shivam H. Patel, Feng Yue, Shannon K. Saw, Rachel Foguth, Jason R. Cannon, Jonathan H. Shannahan, Shihuan Kuang, Arman Sabbaghi, Chad C. Carroll

**Author notes:** Address for correspondence: Chad C. Carroll, PhD Assistant Professor Purdue University, Department of Health and Kinesiology 800 W. Stadium Ave, West Lafayette, IN 47907.

## Abstract

Debilitating cases of tendon pain and degeneration affect the majority of diabetic individuals. The high rate of tendon degeneration persists even when glucose levels are well controlled, suggesting that other mechanisms may drive tendon degeneration in diabetic patients. The purpose of this study was to investigate the impact of advanced glycation end-products on tendon fibroblasts to further our mechanistic understanding of the development and progression of diabetic tendinopathy. We proposed that advanced glycation end-products would induce limitations to mitochondrial function and proliferative capacity in tendon-derived fibroblasts, restricting their ability to maintain biosynthesis of tendon extracellular matrix. Using an *in-vitro* cell culture system, rat Achilles tendon fibroblasts were treated with glycolaldehyde-derived advanced glycation end-products (0, 50, 100, and 200μg/ml) for 48 hours in normal glucose (5.5mM) and high glucose (25mM) conditions. We demonstrate that tendon fibroblasts treated with advanced glycation end-products display reduced ATP production, electron transport efficiency, and proliferative capacity. These impairments were coupled with alterations in mitochondrial DNA content and expression of genes associated with extracellular matrix remodeling, mitochondrial energy metabolism, and apoptosis. Our findings suggest that advanced glycation end-products disrupt tendon fibroblast homeostasis and may be involved in the development and progression of diabetic tendinopathy.

## Introduction

Over 30 million Americans suffer from diabetes. Evaluation of this patient population suggests that tendon abnormalities (i.e., tendinopathies) such as collagen fibril disorganization, increased tissue stiffness, and tissue degeneration are present in the majority of diabetic individuals^1–3^. Notably, the high rate of tendinopathies in diabetic individuals persists even when glucose levels are well controlled (hemoglobin A1c<7)^2, 4, 5^. Given the high prevalence of diabetic tendinopathy, there is a critical need to define the molecular process underlying the diabetic tendon phenotype.

Advanced glycation end-product (AGE) formation is a non-enzymatic process in which free amine terminals are subjected to covalent modification by reactive glucose or other carbonyl containing molecules. AGEs form at an accelerated rate in diabetes and these modifications can result in non-enzymatic cross-links in long-lived extracellular proteins such as tendon collagen, which increase tissue stiffness and alter tissue mechanical properties^6–10^. Furthermore, circulating AGE adducts are able to interact with the receptor for advanced glycation end-products (RAGE) to initiate a noxious feed forward cycle of sustained inflammation and tissue damage^11^. In other tissues, endogenous AGE formation as result of chronic hyperglycemia and spontaneous oxidation of glycolytic intermediates contribute significantly to elevated levels of bound and circulating AGE adducts in diabetic patients^12, 13^. Additionally, AGE-rich diets can increase serum AGE levels and result in accumulation of AGEs in tendon of mice^14^. It is not known, however, what role circulating AGEs play in the development and progression of diabetic tendinopathy.

In non-tendon models, AGEs have been shown to activate RAGE-mediated cellular pathways leading to impairments in mitochondrial function and apoptosis^15–18^. To our knowledge, the impact of AGEs on tenocyte mitochondrial function (e.g., ATP production and electron transport efficiency) has not previously been evaluated. Specifically, ATP has been shown to promote biosynthesis of the extracellular matrix (ECM) in intervertebral cells^19^. In diabetic tendinopathy, degeneration and loss of organization in the ECM may be driven, in part, by the imbalance of energy (ATP) demand and supply. We propose that AGEs contribute to the diabetic tendon phenotype by activating cellular pathways that limit mitochondrial function, thereby interfering with the capacity of tendon fibroblasts to maintain biosynthesis of tendon ECM.

In effort to better understand diabetes associated tendon pathology, we sought to characterize impairments to energy producing systems and proliferative capacity of tendon-derived fibroblasts in response to AGE treatment and high glucose containing medium. We hypothesized that AGEs, *in-vitro*, would impair mitochondrial function and proliferative capacity, independent of glucose concentrations. Further, we hypothesized that AGEs and high glucose medium would reduce mitochondrial DNA (mtDNA) content and further contribute to limitations imposed to energy producing pathways in tendon fibroblasts. To advance our understanding of mitochondrial biogenesis and energy production during AGE exposure, analysis of mitochondrial apoptotic regulators, as well as regulators of mitochondrial energy metabolism was completed. This study provides new functional and descriptive perspective of the AGE insult on tendon fibroblast homeostasis.

## Results

### Proliferative Capacity

Cell proliferation (EdU), cell counts, Mybl2 mRNA, Pcna mRNA, and cytostatic MTT were significantly reduced at AGE-50μg/ml, AGE- 100μg/ml, and AGE-200μg/ml compared to AGE-0μg/ml (p<0.0125, Figure 1b-f, respectively). The HG condition had no effect on cell proliferation (EdU), cell counts, and Mybl2 mRNA (p>0.0125, Figure 1b-d, respectively). However, the HG condition reduced Pcna mRNA transcript counts (p<0.0125, Figure 1e) and increased absorbance values of cytostatic MTT data (p<0.0125, Figure 1f).

**Figure 1:**
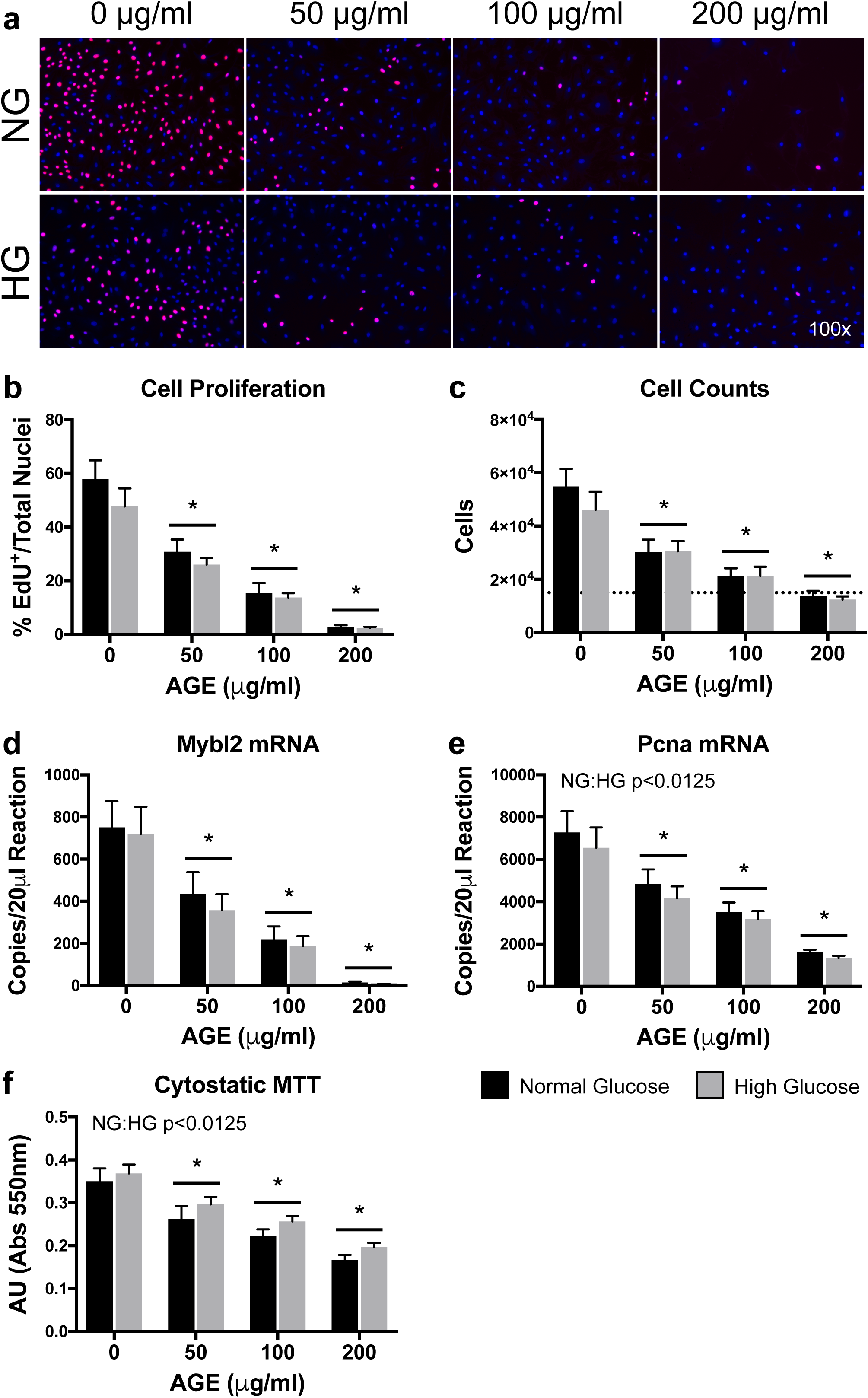
Proliferative Capacity (n=8). **a.** Representative images of EdU^+^/DAPI overlay after 48 hours of AGE treatment. EdU^+^ nuclei are pink and have active DNA synthesis. Blue nuclei (DAPI) do not have active DNA synthesis. **b.** Graphical representation of EdU images (Panel A). Data represented as ratio of EdU^+^ nuclei to total nuclei. **c.** Cell counts after 48 hours of AGE treatment. Dashed line represents initial seeding density of 15,000 cells. **d&e.** ddPCR gene transcript counts for Myb-related protein B (Mybl2) and proliferating cell nuclear antigen (Pcna). **f.** Cytostatic MTT shown as absolute arbitrary absorbance units. *p<0.0125 main effect for AGE vs. AGE-0μg/ml, mixed effects regression. NG:HG p<0.0125 indicates main effect for glucose condition, mixed effects regression. Data presented as mean ± standard error. ■ Normal Glucose (NG), ■ High Glucose (HG).

### Mitochondrial Stress Tests

ATP production-coupled respiration was significantly reduced at AGE-100μg/ml and AGE-200μg/ml compared to AGE-0μg/ml (p<0.0125, Figure 2a). ATP production-coupled respiration was also significantly reduced in the HG condition (p<0.0125, Figure 2a). Basal respiration was significantly reduced at AGE- 100μg/ml and AGE-200μg/ml compared to AGE-0μg/ml (p<0.0125, Figure 2b), but no glucose effect was observed (p>0.0125, Figure 2b). Neither AGE treatment nor glucose condition had a significant effect on maximal respiration (p>0.0125, Figure 2c). AGE treatment had no effect on proton leak (p>0.0125, Figure 2d), but the HG condition increased proton leak across the inner mitochondrial membrane (p<0.0125, Figure 2d). Coupling efficiency was significantly reduced at AGE-200μg/ml compared to AGE- 0μg/ml (p<0.0125, Figure 2e). Coupling efficiency was also significantly reduced in the HG condition (p<0.0125, Figure 2e). Spare respiratory capacity was significantly increased at AGE-200μg/ml compared to AGE-0μg/ml (p<0.0125, Figure 2f), but no glucose effect was observed (p>0.0125, Figure 2f).

**Figure 2:**
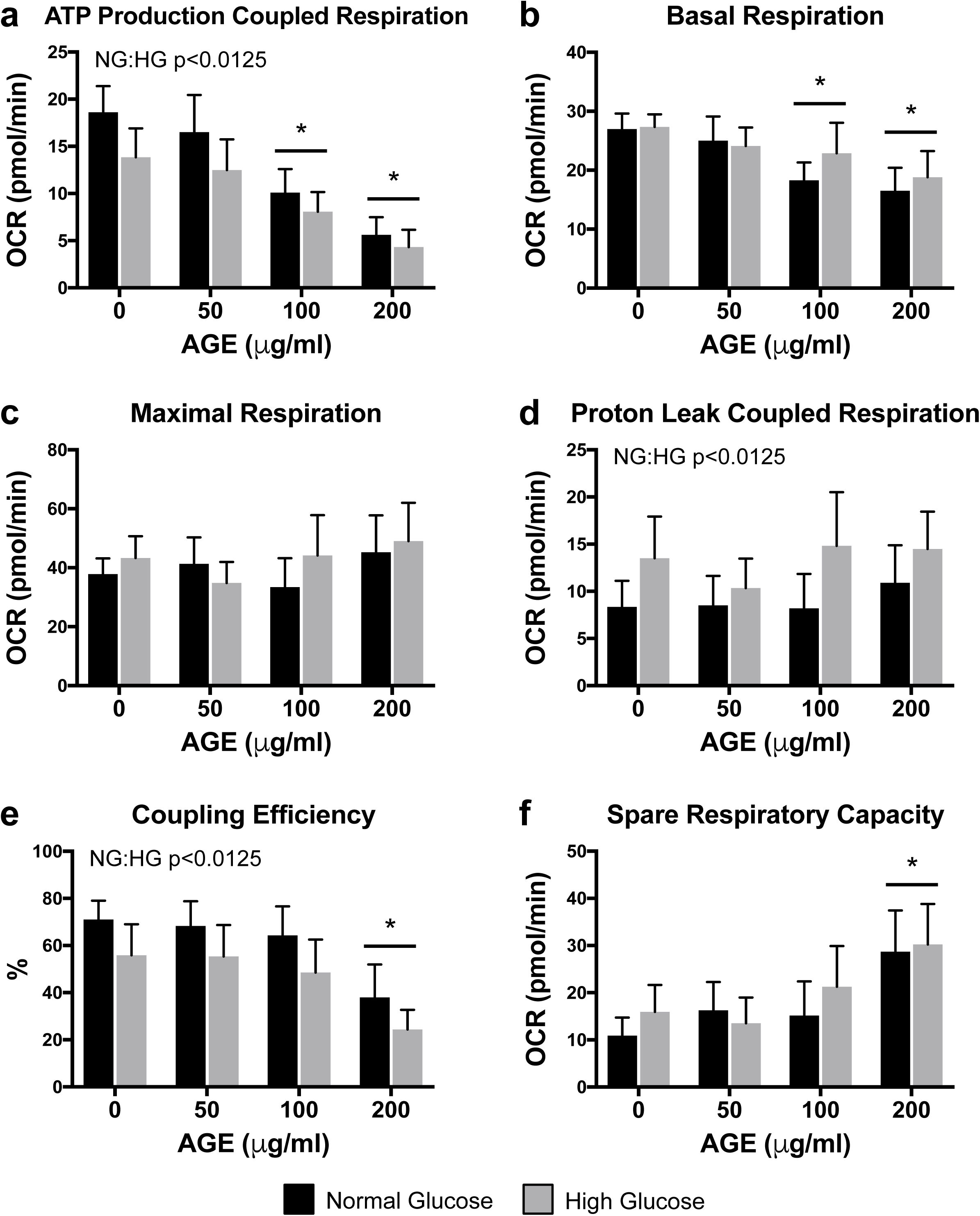
Mitochondrial Stress Tests (n=8). **a-f.** Mitochondrial parameters as a function of oxygen consumption rate (OCR). *p<0.0125 main effect for AGE vs. AGE-0μg/ml, mixed effects regression. NG:HG p<0.0125 indicates main effect for glucose condition, mixed effects regression. Data presented as mean ± standard error. ■ Normal Glucose (NG), ■ High Glucose (HG).

### Transcriptional Analysis of Extracellular Matrix Remodeling

Col1a1 and MMP9 mRNA transcript counts were significantly reduced in the AGE-200μg/ml compared to AGE-0μg/ml (p<0.0125, Figure 3a&d). Col3a1 mRNA transcript counts were significantly increased in the AGE-50μg/ml and AGE-100μg/ml groups compared to AGE-0μg/ml (p<0.0125, Figure 3b). MMP2 mRNA transcript counts were significantly increased in the AGE-100μg/ml compared to AGE-0μg/ml (p<0.0125, Figure 3c). AGEs had no effect on TIMP1 mRNA transcript counts (p<0.0125, Figure 3e). No glucose effect was observed for Col1a1, Col3a1, MMP2, MMP9, or TIMP1 (0<0.0125, Figure 3a-e).

**Figure 3:**
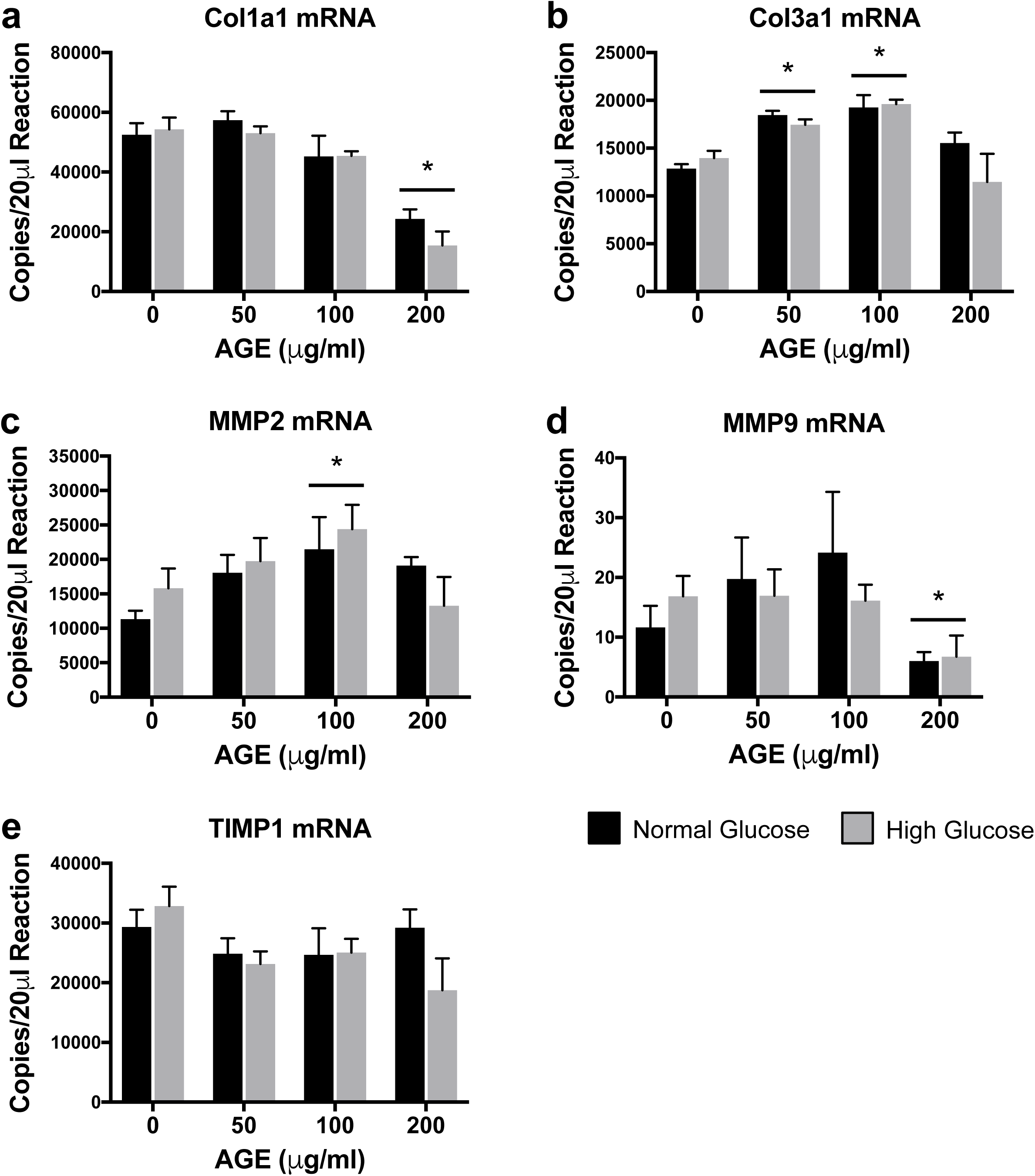
Transcriptional Analysis of Extracellular Matrix Remodeling (n=4). **a-e.** ddPCR gene transcript counts for Collagen alpha-1(I) chain (Col1a1), Collagen alpha-1(III) chain (Col3a1), Matrix metallopeptidase 2 (MMP2), Matrix metalloproteinase 9 (Col3a1), and Tissue inhibitor of matrix metalloproteinase 1 (TIMP1).*p<0.0125 main effect for AGE vs. AGE-0μg/ml, mixed effects regression. Data presented as mean ± standard error. ■ Normal Glucose (NG), ■ High Glucose (HG).

### Mitochondrial DNA Content

mtDNA content was significantly increased at AGE-50μg/ml, AGE-100μg/ml, and AGE-200μg/ml compared to AGE-0μg/ml (p<0.0125, Figure 4a), but no effect of glucose condition was observed (p>0.0125, Figure 4a). Neither AGE treatment nor glucose condition had a significant effect on DNA content when normalized to cell counts (p>0.0125, Figure 4b).

**Figure 4:**
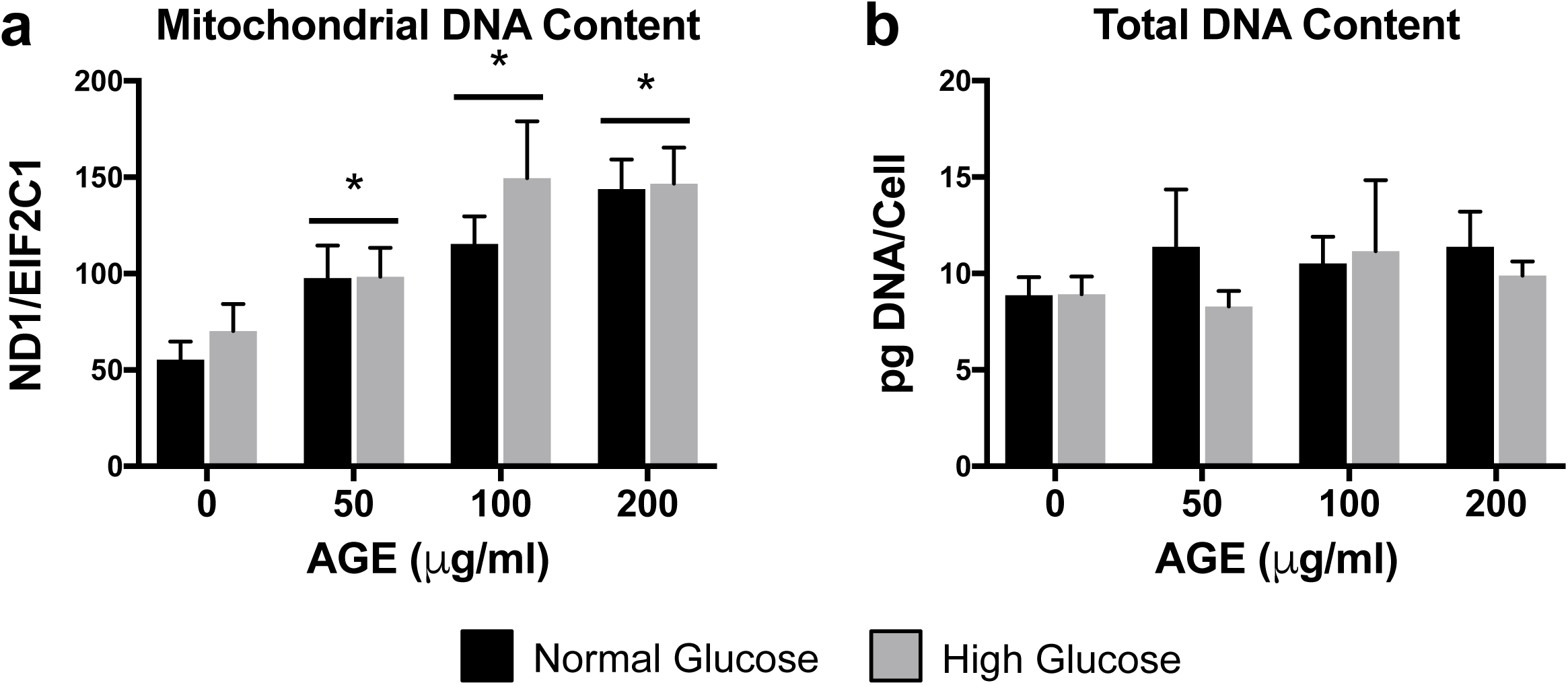
Mitochondrial DNA Content (n=8). **a.** mtDNA content quantified by ratio of mitochondrial gene, NADH dehydrogenase subunit 1 (ND1), to nuclear reference gene, protein argonaute-1 (EIF2C1). **b.** Total DNA content normalized to cell counts. *p<0.0125 main effect for AGE vs. AGE-0μg/ml, mixed effects regression. Data presented as mean ± standard error. ■ Normal Glucose (NG), ■ High Glucose (HG).

### Transcriptional Analysis of Mitochondrial Complexes

Ndufa1 mRNA transcript counts were significantly increased at AGE-50μg/ml, AGE-100μg/ml, and AGE-200μg/ml compared to AGE-0μg/ml (p<0.0125, Figure 5a), but no effect of glucose condition was observed (p>0.0125, Figure 5a). Sdha mRNA transcript counts were significantly reduced at AGE-200μg/ml compared to AGE-0μg/ml (p<0.0125, Figure 5b), but no effect of glucose was observed (p>0.0125, Figure 5b). A secondary direct comparison to test the conditional main effect revealed that Sdha transcript counts were significantly reduced at HG AGE-200μg/ml compared to HG AGE-0μg/ml (p<0.05, Figure 5b). Bcs1l mRNA was significantly reduced at AGE-50μg/ml, AGE-100μg/ml, and AGE-200μg/ml compared to AGE-0μg/ml (p<0.0125, Figure 5c). Bcs1l mRNA was also significantly reduced in the HG condition (p<0.0125, Figure 5c). Neither AGE treatment nor glucose condition altered Cox4i1 mRNA transcript counts (p>0.0125, Figure 5d). MT-ATP6 mRNA transcript counts were significantly reduced at HG AGE-200μg/ml compared to HG AGE-0μg/ml (p<0.05, Figure 5e), but no effect of glucose was observed (p>0.0125, Figure 5e).

**Figure 5:**
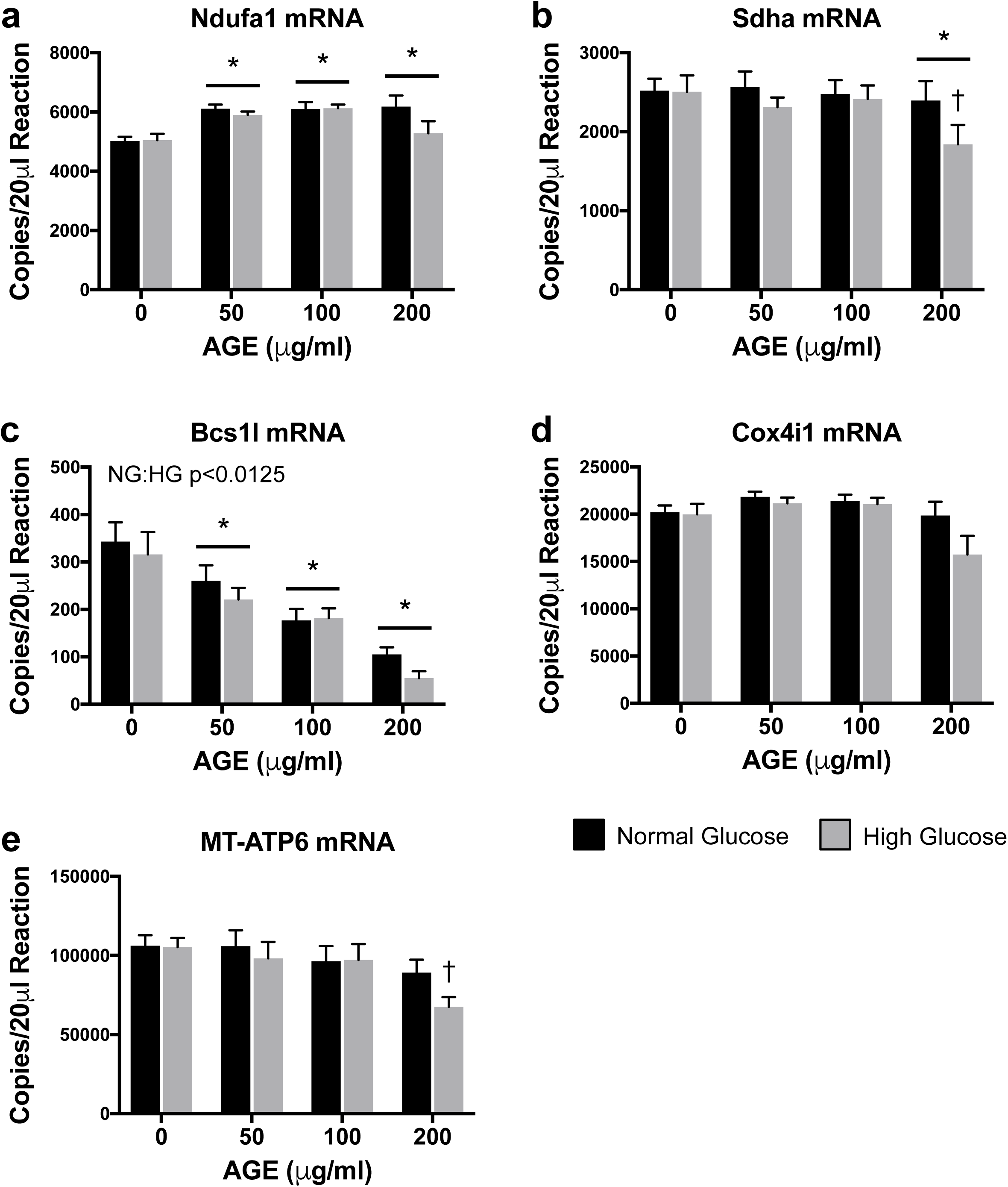
Transcriptional Analysis of Mitochondrial Complexes (n=8). **a-e.** ddPCR gene transcript counts for NADH:ubiquinone oxidoreductase subunit A1 (Ndufa1), Succinate dehydrogenase complex flavoprotein subunit A (Sdha), BCS1 homolog, ubiquinol-cytochrome c reductase complex chaperone (Bcs1l), Cytochrome c oxidase subunit 4i1 (Cox4i1), and ATP synthase 6, mitochondrial (MT-ATP6). *p<0.0125 main effect for AGE vs. AGE-0μg/ml, mixed effects regression. †p<0.05 vs. HG AGE-0μg/ml, paired t-test. NG:HG p<0.0125 indicates main effect for glucose condition, mixed effects regression. Data presented as mean ± standard error. ■ Normal Glucose (NG), ■ High Glucose (HG).

### Protein Analysis of Mitochondrial Complexes

Expression of complex III (UQCRC2) was significantly increased at AGE-200μg/ml compared to AGE-0μg/ml (p<0.017, Figure 6c), but no glucose effect was observed (p>0.017, Figure 6c). Expression of complex I (NDUFB8), II (SDHB), IV (MTCO1), or V (ATP5A) were not altered by either AGE treatment or glucose condition (p>0.017, Figure 6a, b, d, & e, respectively). However, a significant interaction between AGE treatment and glucose condition was observed for complex I (NDUFB8) and complex IV (MTCO1) (p<0.017, Figure 6a&d).

**Figure 6:**
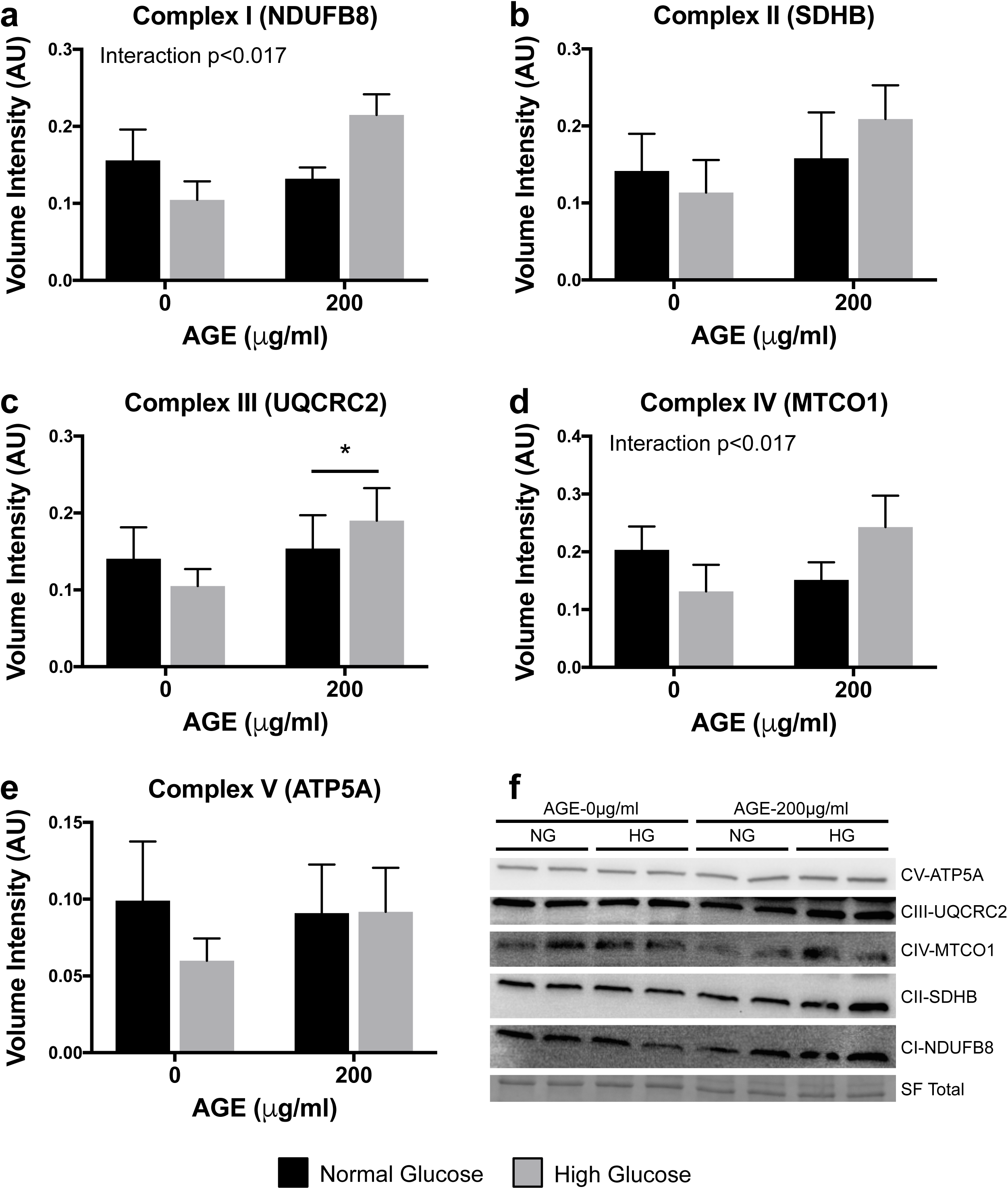
Protein Analysis of Mitochondrial Complexes (n=4). **a-e**. OXPHOS protein expression for mitochondrial complex I (NDUFB8), complex II (SDHB), complex III (UQCRC2), complex IV (MTCO1), and complex V (ATP5A). F. Representative blot images (displayed in order of molecular weight) including Stain Free (SF) UV visualized total protein. All targets were probed on the same membrane, but are cropped due to different exposure times required for each target (Signal Accumulation Mode). *p<0.017 main effect for AGE vs. AGE-0μg/ml, mixed effects regression. Interaction p<0.017, significant interaction between AGE treatment and glucose condition. Data presented as mean ± standard error. ■ Normal Glucose (NG), ■ High Glucose (HG).

### Transcriptional Analysis of Mitochondrial Apoptosis

Bak1 mRNA transcript counts were significantly reduced at AGE-100μg/ml and AGE-200μg/ml compared to AGE-0μg/ml (p<0.0125, Figure 7a), but no effect of glucose condition was observed (p>0.0125, Figure 7a). Bax and Casp8 mRNA transcript counts were significantly reduced at HG AGE-200μg/ml compared to HG AGE-0μg/ml (p<0.05, Figure 7b&e, respectively), but no effect of glucose was observed (p>0.0125, Figure 7b&e, respectively). Bcl2 and Tgfbr3 mRNA transcript counts were significantly reduced at AGE-200μg/ml compared to AGE-0μg/ml (p<0.0125, Figure 7c&f, respectively), but no effect of glucose condition was observed (p>0.0125, Figure 7c&f, respectively). AGE treatment had no effect on Casp3 mRNA (p>0.0125, Figure 7d), but the HG condition reduced Casp3 mRNA transcript counts (p<0.0125, Figure 7d).

**Figure 7:**
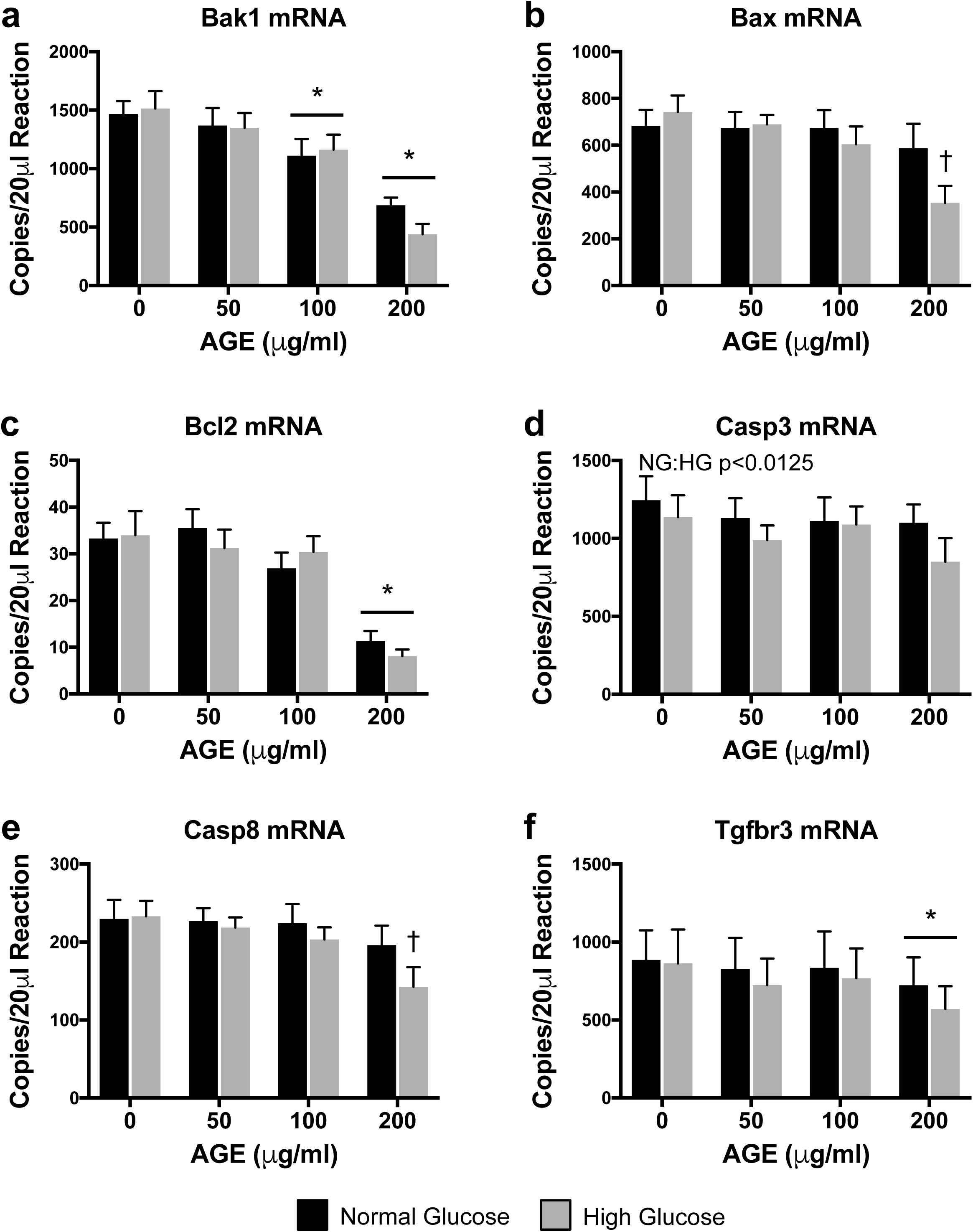
Transcriptional Analysis of Mitochondrial Apoptosis (n=8). **a-f.** ddPCR gene transcript counts for BCL2-antagonist/killer 1 (Bak1), BCL2 associated X, apoptosis regulator (Bax), B-cell lymphoma 2, apoptosis regulator (Bcl2), Caspase 3 (Casp3), Caspase 8 (Casp8), and Transforming growth factor beta receptor 3 (Tgfbr3). *p<0.0125 main effect for AGE vs. AGE-0μg/ml, mixed effects regression. †p<0.05 vs. HG AGE-0μg/ml, paired t-test. NG:HG p<0.0125 indicates main effect for glucose condition, mixed effects regression. Data presented as mean ± standard error. ■ Normal Glucose (NG), ■ High Glucose (HG).

### Superoxide Production

Superoxide production was significantly increased at AGE-200μg/ml compared to AGE-0μg/ml (p<0.017, Figure 8), but no glucose effect was observed (p>0.017, Figure 8).

**Figure 8:**
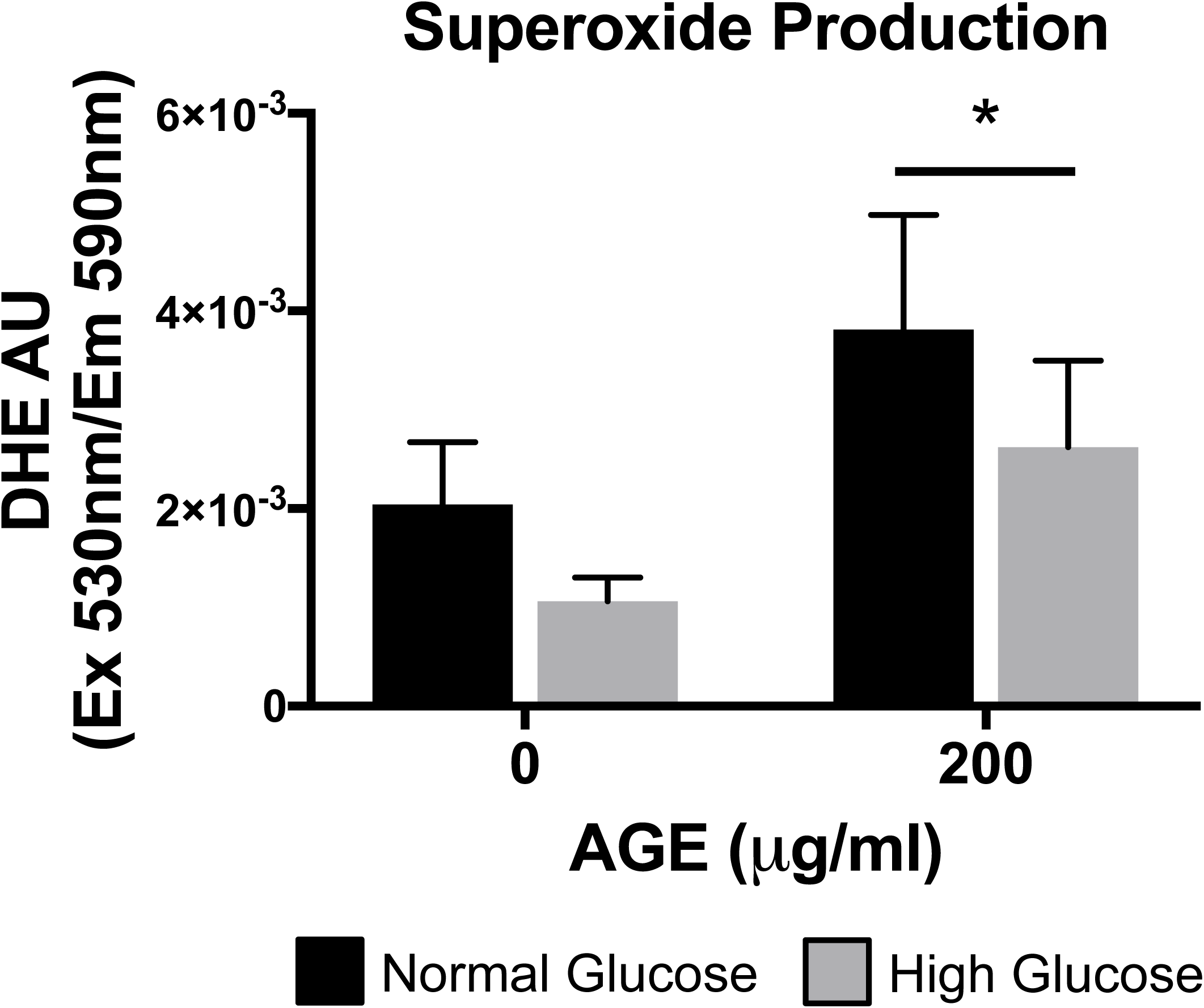
Superoxide Production (n=4). *p<0.017 main effect for AGE vs. AGE-0μg/ml, mixed effects regression. Data presented as mean ± standard error. ■ Normal Glucose (NG), ■ High Glucose (HG).

## Discussion

AGE-induced non-enzymatic collagen cross-link formation in the ECM has been proposed to increase tendon stiffness in diabetic individuals^6, 7, 10^. However, factors contributing to tendon degeneration and collagen fibril disorganization in diabetic tendon have not been extensively characterized. The detrimental effects of AGE associated pathology are well described in non-tendon tissues and have been linked to several diabetic complications such as cardiomyopathy, retinopathy, nephropathy, and endothelial dysfunction^10, 20^. In this study, we sought to address the effect of AGEs at the cellular level to better understand factors that may contribute to the disorganization and degeneration of tendon ECM noted in diabetic individuals. Using an *in-vitro* primary cell culture system, we demonstrate dose-dependent AGE-induced reductions in proliferative capacity and mitochondrial ATP production of tendon-derived fibroblasts. Additionally, we demonstrate increased mtDNA content and modifications to mitochondrial complexes and markers of apoptosis after AGE treatment. While previous research has established AGE dependent mitochondrial and proliferative limitations^17, 18, 21^, these data, to the best of our knowledge, are the first to show these impairments in tendon-derived fibroblasts.

Tendon injuries and degenerative pathology are a common and debilitating clinical problem in diabetic individuals^2, 3, 22–26^. Diabetic tendons are thicker^24, 27^, and more commonly present with fibril disorganization^2^. Despite convincing epidemiological data, the molecular factors contributing to the development and degenerative process of tendinopathy in diabetic individuals are not well characterized. Much focus has been directed to the influence of elevated glucose on cellular and structural tendon parameters. *In-vitro* and *in-vivo* data has indicated that elevated glucose availability can alter cell signaling dynamics and structural properties in tendon^28–30^. These data suggest that glucose may contribute to tendon pathology, however conflicting reports exist and more conclusive human data is needed to confirm hyperglycemia-associated tendon degeneration in diabetes.

As evidenced, glucose does seem to be implicated in modulation of some tendon cellular and structural properties^28–30^, however, the underlying mechanisms influencing tendon degeneration in diabetic patients remain inconclusive. Couppé *et al.*^4^ have demonstrated that Achilles tendon Young’s modulus (stiffness) is greater in diabetic subjects compared to control subjects, but no difference exists between subjects with well-controlled and poorly controlled diabetes. These data highlight that hyperglycemia is unlikely to be the sole contributor of diabetic tendon pathology. In support of this notion, we explored the role of AGEs on modulation of tendon cellular properties that may consequently interfere with regulation and maintenance of the ECM.

Tendon fibroblast proliferation is a vital component to tendon development, adaptation, and healing^31, 32^. An inability of tenocytes to proliferate in the presence of AGEs would significantly precipitate the development of tendinopathy^38^. Tendinopathy is thought to develop, in part, from a failed healing response to minor tendon damage during loading. Specifically, delayed and abnormal healing is a common complication of both type I and type II diabetes^33, 34^. As evidence, tendinopathies in diabetic patients are generally more pervasive and can present with more severe degeneration, which may, in part, be driven by prolonged injury status^2, 35, 36^. We note dramatic impairments to tendon fibroblast proliferative capacity in the presence of AGEs. Our *in-vitro* data demonstrate, in a dose dependent fashion, reduced cell proliferation (EdU) and cell counts after 48 hours of AGE treatment (Figure 1b&c, respectively). In concert, we demonstrate a reduction in proliferative gene markers, Mybl2 and Pcna, and reduced absorbance values of cytostatic MTT with AGE treatment (Figure 1d-f, respectively). These data are in agreement with data from Hu *et al*.^21^, which also demonstrates a lack of proliferative capacity in the presence of AGEs. While the HG condition did impact Pcna mRNA and cytostatic MTT (Figure 1e-f), the primary insult to proliferative capacity appears to be driven by AGE treatment. Impaired proliferation is likely mediated by RAGE signaling^37^, but further work is needed to confirm AGE-mediated impairments to tendon fibroblast proliferation and tendon healing.

AGEs have been linked to numerous diabetes related complications and their detrimental effects are well documented in several tissues^10, 20^. The primary consequences of AGE-mediated RAGE activation are chronic inflammation, as result of the NF-κB mediated cascade, and oxidative stress, due to NADPH oxidase stimulation. Among resulting complications, reports in other cell types have identified AGE-mediated impairments to mitochondrial function and dynamics^15–18^. Similarly, we report impaired mitochondrial parameters and reduced ATP production-coupled respiration in tendon fibroblasts treated with AGEs (Figure 2a). The role of ATP is diverse and essential for a multitude of cellular processes including cell proliferation, contraction, and wound healing. Specifically, ATP has been shown to promote biosynthesis of the ECM in intervertebral disk cells^19^. In tendon, the resident fibroblast population maintains the ECM, which is primarily composed of collagen. Due to the energy demanding nature of ECM maintenance, it is possible, that in the presence of AGEs, limited ATP production contributes to loss of organization in the ECM, which is commonly noted in diabetic tendon^1–3^. While maximal respiration of tendon fibroblasts was unchanged due to identical XFp seeding density, basal respiration was reduced after AGE treatment (Figure 2b&c, respectively). In addition to loss of ATP production and reduced basal respiration, we demonstrate that AGEs also impair electron transport coupling efficiency; thereby increasing spare respiratory capacity of AGE treated tendon fibroblasts and indicating overall reduction to electron transport efficiency (Figure 2e&f, respectively). Interestingly, the HG condition increased proton leak across the inner mitochondrial membrane and reduced ATP production and coupling efficiency, suggesting that glucose alone can also impact electron transport efficiency (Figure 2d). It is important to note that AGE and/or glucose-mediated mitochondrial impairments are likely not the sole contributor of tendon ECM disorganization. For example, AGEs increase matrix metalloproteinases (MMP) expression and activity, which may contribute in an additive manner to enhanced degeneration in tendon ECM^38^.

To address our hypothesis that the presence of AGEs and resulting limited ATP production contribute to loss of organization in the ECM, we completed gene analysis of targets associated with ECM maintenance and remodeling. We note that AGE treatment reduces Col1a1 mRNA expression (Figure 3a), consistent with previous work which has demonstrated reduced collagen synthesis during AGE treatment in fibroblasts^39^. Col3a1 mRNA expression was however increased with AGE 50μg/ml and 100μg/ml, but not at 200μg/ml (Figure 3b). Col3a1 expression is upregulated after injury and during the early stages of wound healing^40, 41^. It is possible that in our cell culture model, increased Col3a1 mRNA expression in the AGE 50μg/ml and 100μg/ml groups was an attempt to overcome the AGE insult to tendon fibroblast collagen production. However, it appears that the potential compensatory response was unsuccessful in the AGE 200μg/ml group (Figure 3b). Further, MMPs and tissue inhibitors of metalloproteinases (TIMPs) work to tightly regulate remodeling of ECM remodeling. Previous work in chondrocyte cell culture has demonstrated an increase to MMP mRNA expression with AGE treatment^42, 43^. While we did observe increased MMP2 mRNA expression in the AGE 100μg/ml group, MMP9 mRNA expression was reduced in the AGE 200μg/ml group (Figure 3c&d). Lastly, we did not observe an AGE or glucose effect of TIMP1 mRNA (Figure 3e. Reduced collagen expression and altered MMP modulation may favor an environment that promotes loss of ECM organization. Future work evaluating post-translational MMP activation and collagen fibril organization is needed to determine the role of AGEs in modulation of tendon ECM.

We also noted increased content of mtDNA after 48 hours of AGE treatment, despite striking reductions to ATP production. mtDNA content can be used as an estimate of mitochondrial volume^44^, however, contrary to our hypothesis, 48 hours of AGE treatment resulted in increased mtDNA content in a dose dependent manner independent of glucose condition (Figure 4a). Based on our findings, it is plausible that the increase in mtDNA in AGE-treated tendon fibroblasts was indicative of a compensatory response to overcome the AGE insult and meet energy demands^45, 46^. In support of this theory, cells under oxidative stress have been shown to increase mitochondria and mtDNA^47^. Additionally, Kim *et al*.^46^ have demonstrated a relationship between mtDNA content and severity of histopathology in cancerous lesions, suggesting that increased mtDNA content may be used as a measure of DNA injury and pathology. Despite this compensatory response, functional mitochondrial limitations were still evident after 48 hours of AGE treatment. Alternatively, it is possible that AGE-induced limitations to mitochondrial biogenesis may result in failure of tendon fibroblasts to meet energy demands. For example, mtDNA is more susceptible to mutation than nuclear DNA and AGEs have been shown to induce damage to mtDNA, which could ultimately impact mitochondrial biogenesis^48, 49^. However, further molecular work is needed to conclusively define the pathway of AGE-mediated mitochondrial damage. To confirm that mtDNA content measurements were not falsely elevated by an increase in total DNA, despite equal DNA loading per PCR reaction, we normalized total DNA yield to cell counts and noted no difference in DNA content with either glucose condition or AGE treatment (Figure 4b).

Reactive oxygen species (ROS), produced primarily by mitochondria, have been implicated in AGE-mediated cell apoptosis and mitochondrial damage^18, 50, 51^. To further explore and identify specific targets of the AGE insult, we assessed regulation of electron transport complexes and apoptosis. Analysis of gene transcripts associated with complex I-V (Figure 5a-e, respectively) of the electron transport chain revealed increased expression of Ndufa1 mRNA (Figure 5a) and marked reduction of Bcs1l mRNA (Figure 5c) with AGE treatment. Additionally, the HG condition further reduced Bcs1l mRNA expression (Figure 5c). Increased Ndufa1 mRNA after AGE treatment may be interpreted as a compensatory response to increase mitochondrial complex I in effort to meet energy demands after the AGE insult. However, Bcs1l mRNA, which encodes mitochondrial complex III, is dramatically reduced in a dose dependent manner after AGE treatment, suggesting that the AGE insult may be targeted to complex III. Lastly, Sdha and MT-ATP6 mRNA, coding for mitochondrial complex II and complex V (ATP synthase), respectively, were reduced in the HG AGE-200μg/ml group compared to HG AGE-0μg/ml (Figure 5b&e, respectively). A direct comparison was made between HG AGE-0μg/ml and HG AGE-200μg/ml to test the conditional main effect and reveal a significant reduction to Sdha and MT-ATP6 mRNA, suggesting that both AGE treatment and HG condition were contributing to this reduction. Cox4i1 (complex IV, Figure 5d) was not impacted by either AGE treatment or HG condition. While these gene data provide indication that AGEs may indeed have targeted effects to the electron transport chain, protein analysis revealed only complex III (UQCRC2) to be affected by an AGE main effect (Figure 6c). Additionally, complex III protein expression was increased with AGE exposure, while complex III mRNA expression was reduced. Lack of response to remaining complexes and discrepancies between gene and protein analysis may be attributed to the acute duration of AGE exposure. Further, an interaction between AGEs and glucose condition was observed in complex I (NDUFB8, Figure 6a) and complex IV (MTCO1, Figure 6d). These results merit further work to discover mechanistic pathways by which AGEs limit mitochondrial function during long-term AGE exposure.

To assess the impact of AGEs on mitochondrial-related tendon fibroblast apoptosis, we completed analysis of associated gene transcripts and measurement of superoxide production. Transcript counts of pro-apoptotic gene Bak1 were reduced with AGE treatment (Figure 7a). Additionally, pro-apoptotic Bax mRNA expression was reduced only in the HG AGE-200μg/ml group compared to HG AGE-0μg/ml (Figure 7b). Although these findings are contrary to previous work^21^, it is possible that these may be compensatory responses to AGE-mediated apoptotic signals. In agreement with previous work by Hu *et al*.^21^, expression of Bcl2 mRNA, an anti-apoptotic mediator, was reduced with AGE treatment (Figure 7c). Previous reports have indicated an inverse relationship between Bcl2 and ROS, where ROS may act to reduce expression of Bcl2 and sensitize cells to apoptosis^52^. In agreement, our data demonstrate increased production of superoxide, a reactive anion species, after AGE treatment (Figure 8). Further, cytochrome c release is suppressed by Bcl2, but is promoted by Bak1 and Bax. Cytochrome c release will ultimately result in caspase activation^53^. Previous reports indicate AGE treatment induces caspase activation^21, 54^, however we did not note any AGE-induced changes to Casp3 mRNA (Figure 7d) but did note reduced Casp8 mRNA in the HG AGE-200μg/ml group compared to HG AGE-0μg/ml (Figure 7e). Finally, transcript counts of Tgfbr3 were reduced with AGE treatment (Figure 7f). Tgfbr3 overexpression has been shown to increase Bax and Bcl2 expression, as well as promote activation of caspase 3^55^. Casp3 mRNA expression was reduced in the HG condition and was the only target associated with apoptosis to be affected by glucose (Figure 7d). From these data, it is evident that AGEs alter transcriptional regulation of gene transcripts associated with apoptosis, however, we are limited in our interpretation and further post-translational and activity assays are needed to define the precise mechanisms by which AGEs may induce apoptosis in diabetic tendinopathy.

In summary (Figure 9), we demonstrate that AGEs, which are elevated in serum of diabetic individuals, impair proliferative capacity, ATP production, and electron transport chain efficiency. Additionally we demonstrate that AGEs alter regulation of mitochondrial complex expression and gene transcripts associated with apoptosis. While the HG condition did impact some mitochondrial parameters, AGEs appear to be the primary insult and may be responsible for the development of the diabetic tendon phenotype. This work provides new insights to the pathophysiology of tendon ECM in diabetic patients. However, future *in-vivo* and mechanistic work is needed to determine whether controlling serum AGEs in diabetic patients can reduce risk of degenerative tendinopathy.

**Figure 9:**
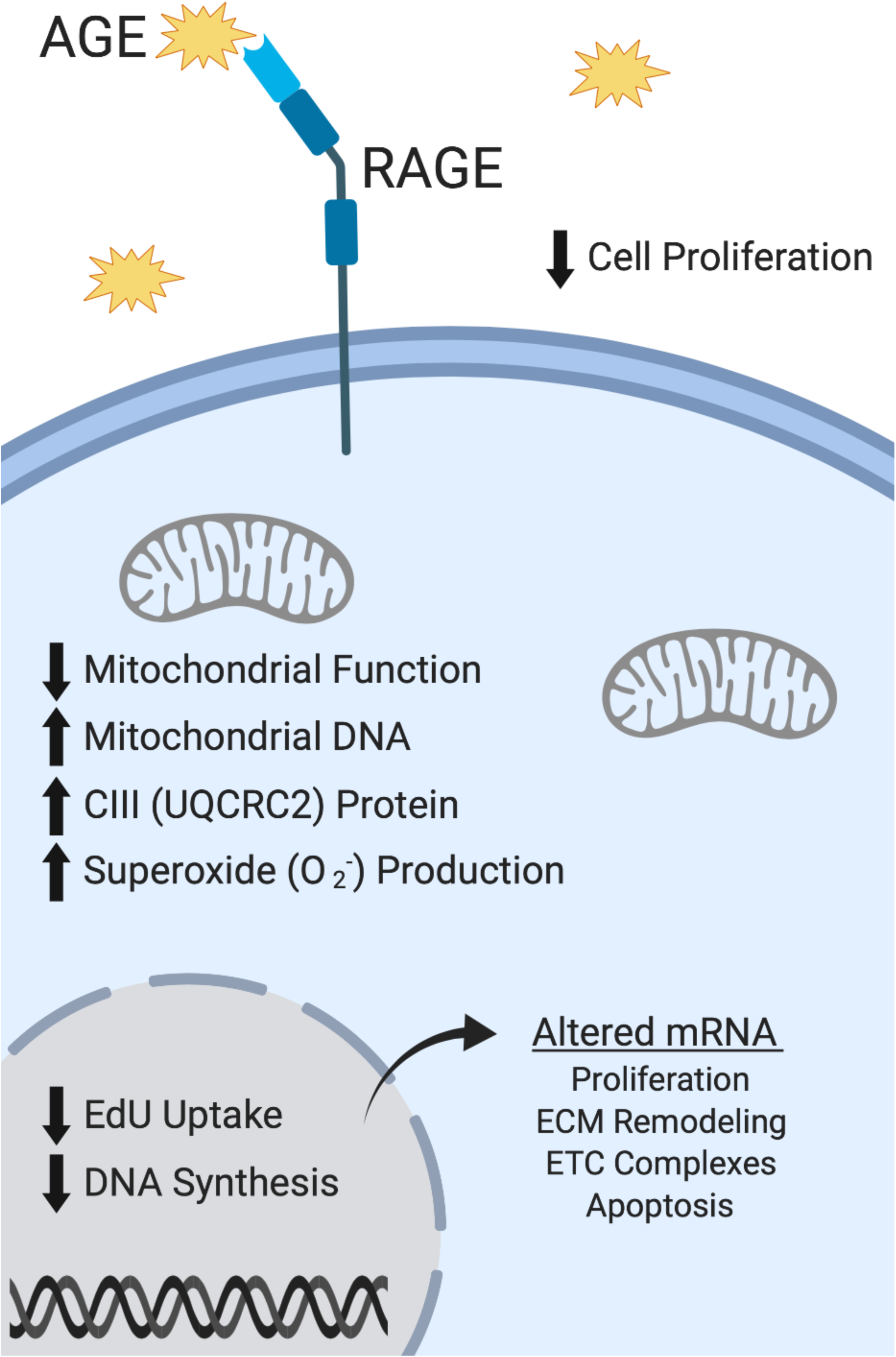
Summary of Major AGE-Mediated Findings. Figure created with BioRender.

## Methods

### AGE Preparation

Glycolaldehyde-derived AGEs were generated under sterile conditions as described by Valencia *et al.*^56^. Briefly, sterile filtered 30% BSA solution (Sigma, St. Louis, MO) was incubated with 70mM glycolaldehyde dimer (Sigma) in sterile PBS without calcium chloride and magnesium chloride for three days at 37°C. After incubation, the AGE product was dialyzed against sterile PBS for 24 hours at 4°C using gamma-irradiated 10kDa cut-off cassettes (Thermo Scientific, Waltham, MA) to remove unreacted glycolaldehyde. Unmodified control BSA was prepared similarly, without the addition of glycolaldehyde dimer. Protein concentration was determined by BCA assay (Thermo Scientific) and absence of endotoxin (<0.25Eu/ml) was confirmed via the LAL gel-clot assay (GenScript, Piscataway, NJ).

### Extent of AGE Modification

Extent of BSA modification was confirmed by fluorescence, absorbance, and loss of primary amines^56–59^. AGE-BSA and Control-BSA were diluted to 1mg/ml in PBS and fluorescent spectra and absorbance were recorded at 335nm excitation/420nm emission and 340nm, respectively (Molecular Devices, San Jose, CA). For determination of loss of primary amines AGE-BSA and Control-BSA were diluted to 0.2mg/ml in PBS. An equal volume of ortho-phthalaldehyde solution (Sigma) was added and fluorescent spectrum was recorded at 340nm excitation/455nm emission (Molecular Devices). A standard curve of 0 to 0.2mg/ml of BSA that did not undergo 37°C incubation was used to generate a standard curve of free amine content and data was normalized to represent percentage of amine terminals remaining with reference to the standard curve^56^. Respective fluorescence or absorbance values of PBS alone were subtracted from all data. AGE-BSA fluorescence was increased to 697.78 arbitrary units (AU) compared to -0.72 AU for control BSA. Absorbance readings were completed to determine the extent of glycation. AGE-BSA showed increased glycation with absorbance readings of 0.682 AU compared to 0.01 AU for control BSA. Finally, AGE-BSA primary amine terminals underwent complete modification (-0.03% accessible amine terminals remaining), while control BSA retained 81.48% of accessible amine terminals. Spectra and absorbance values of PBS alone were subtracted from data but resulted in negative values for fluorescent-based detection because the spectra of PBS alone was greater than the obtained sample values. Negative values were interpreted as zero, and extent of modification was similar to previous reports^56^.

### Animal Protocol

This study was approved by the Purdue University Institutional Animal Care and Use Committee and all animals were cared for in accordance with the recommendations in the Guide for the Care and Use of Laboratory Animals^60^. Eight-week-old female Sprague-Dawley rats (n=8, 256.43±5.19g) were purchased from Charles River Laboratories (Wilmington, MA) and maintained for an additional eight weeks. Rats were housed on a 12-hour light-dark cycle and provided access to standard rat chow and water ad libitum. At sixteen weeks, rats were euthanized by decapitation after CO_2_ inhalation.

### Tendon Fibroblast Isolation and Cell Culture

Tendon-derived fibroblasts were isolated from the Achilles tendons of eight rats. After euthanasia, Achilles tendons were carefully excised and trimmed of remaining muscle and fascia. The tendon was rinsed with sterile PBS, finely minced, placed in DMEM containing 0.2% type I collagenase, and incubated in a 37°C shaking water bath for four hours. After tissue digestion, the cell suspension was filtered through a 100μm mesh filter, pelleted by centrifugation, resuspended in 5.5mM glucose DMEM containing 10% FBS, 1% sodium pyruvate (Sigma), and 1% penicillin/streptomycin (Thermo Scientific) and plated in 100mm collagen coated dishes. After reaching ∼75-80% confluence, tendon fibroblasts were subcultured with either normal glucose (NG, 5.5mM) DMEM (Sigma) or high glucose (HG, 25mM) DMEM (Sigma) for a minimum of one passage to allow for glucose condition acclimation. Tendon fibroblasts were then seeded into 6-well, 24-well, or 96-well collagen coated culture plates and treated for 48 hours with 0, 50, 100, and 200μg/ml of AGEs within both glucose conditions. The 0μg/ml AGE group was treated with 200μg/ml of unmodified control BSA. Tendon fibroblasts from passage 2-4 were used for all experiments.

### Proliferative Capacity

Proliferative capacity was assessed by cell counts, cytostatic MTT, and incorporation of synthetic nucleoside 5-ethynyl-2-deoxyuridine (EdU). After AGE treatment in 6-well plates, cells were enzymatically released and manually counted with a hemocytometer. Cell counts were completed in duplicate and normalized to total volume in which cells were resuspended. For cytostatic MTT analysis, cells cultured in a 96-well plate were incubated with MTT reagent (5mg/ml, VWR, Radnor, PA) for the final 90 minutes of AGE treatment. Formazan crystals were solubilized with DMSO and absorbance was read at 550nm (Multiskan, Thermo Scientific). Each assay was completed in duplicate and data was represented as absolute arbitrary absorbance units. Cells were labeled with 2.5μM EdU (Carbosynth, Newbury, UK) during the final 12 hours of AGE treatment. EdU is a thymidine analog that incorporates during active DNA synthesis and is used to mark proliferating cells. Fixed cells (4% PFA) were stained with DAPI, and EdU was visualized via the Click-iT method (100mM Tris-HCl pH8.5, 1mM CuSO_4_, 2.5μM red-fluorescent tetramethylrhodamine azide, and 100mM ascorbic acid; Thermo Scientific)^61^. Fluorescent images were captured using a DMI 6000B microscope (Leica, Wetzlar, Germany) with a 10x objective and MetaMorph software (Molecular Devices). EdU data was analyzed using the ImageJ Multi-Point Tool (National Institutes of Health, Bethesda, Maryland) and is reported as a ratio of EdU^+^ nuclei to total nuclei from two fields-of-view within each treatment.

### Mitochondrial Stress Tests

Tendon fibroblasts were treated for 48 hours prior to being plated (8,000 cells) into collagen coated XFp cell culture miniplates (Seahorse Bioscience, Agilent, Santa Clara, CA). Cultured cells from each treatment condition were reseeded in duplicate, allowed to attach to the miniplates, and washed twice in pre-warmed XF base medium (Agilent). Miniplates were then incubated in XF base medium (1mM sodium pyruvate, 2mM glutamine, and either 5.5mM or 25mM glucose) for one hour in a 37°C incubator without CO_2_ to allow plate outgassing. Mitochondrial stress experiments were completed on an XFp extracellular flux analyzer (Agilent) with the Mito Stress Test Kit (Agilent). Chemicals (1μM oligomycin, 0.5μM carbonyl cyanide-*4*-(trifluoromethoxy)phenylhydrazone (FCCP), 0.5μM rotenone, and 0.5μM antimycinA) were preloaded into cartridge ports and injected in succession during measurement of oxygen consumption rate (OCR).

### Gene Expression Analysis

Total RNA for gene expression analysis was isolated after AGE treatment using the Direct-zol RNA Miniprep kit (Zymo Research). On-column DNase digestion was completed on all samples prior to elution of RNA. RNA concentration was determined using a NanoDrop 2000 (Thermo Scientific). Quality of RNA was assessed using the 260/280 and 260/230 ratios. Reverse transcription (iScript, BioRad) was completed to produce complementary DNA from 100ng of RNA. Absolute quantification of mRNA target transcripts was completed using the ddPCR platform (BioRad) with validated probe-based assays (BioRad) as previously described^30^. For optimal detection, Col1a1 and Col3a1 reactions contained 0.55ng of cDNA, while MMP2, MMP9, and TIMP1 contained 4.4ng of cDNA^30^. All other target PCR reactions contained 2.2ng of cDNA. A list of measured gene targets is provided in Table 1.

**Table 1:**
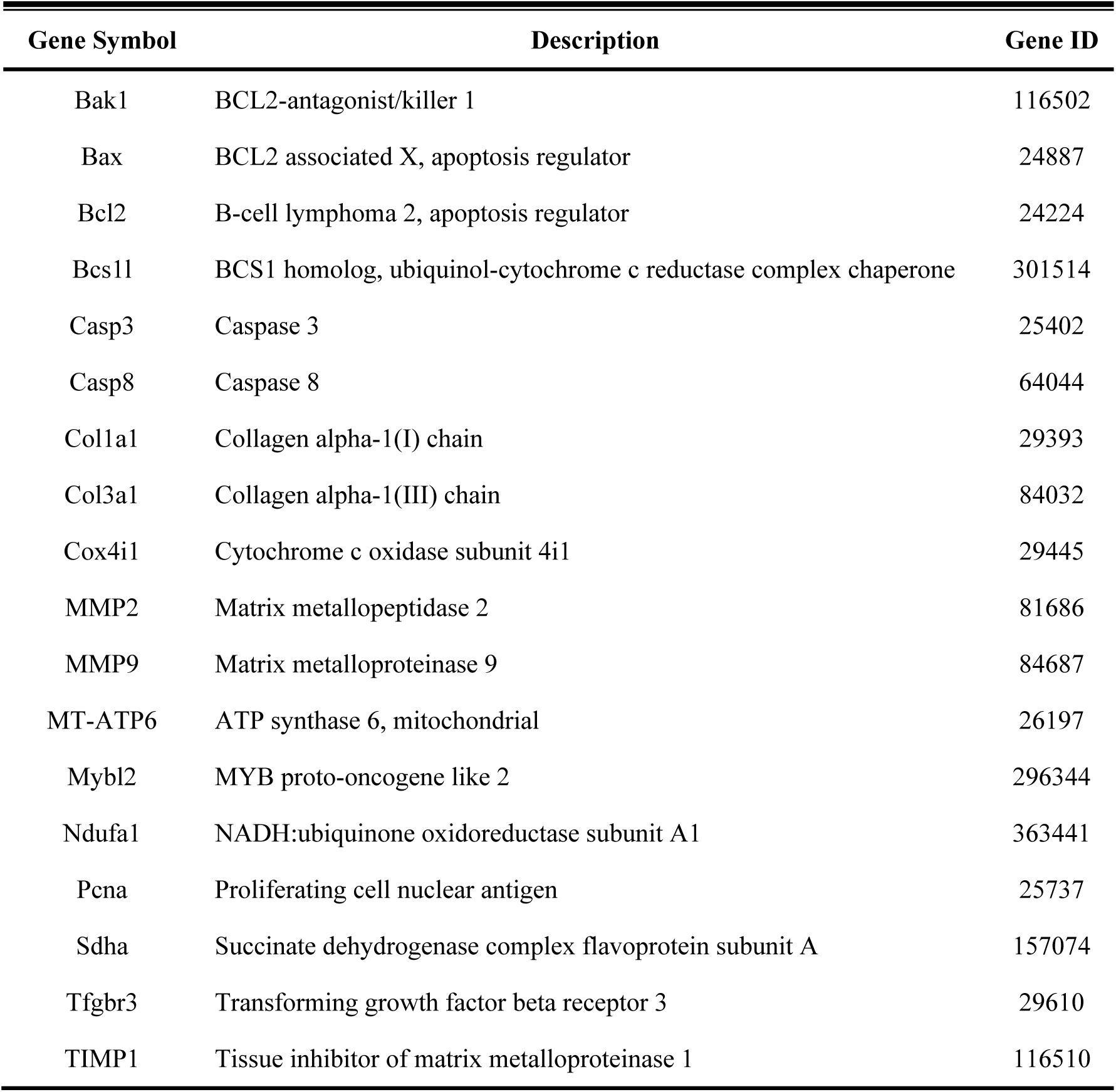
ddPCR Gene Targets

| Gene Symbol | Description | Gene ID |
| --- | --- | --- |
| Bak1 | BCL2-antagonist/killer 1 | 116502 |
| Bax | BCL2 associated X, apoptosis regulator | 24887 |
| Bcl2 | B-cell lymphoma 2, apoptosis regulator | 24224 |
| Bcs1l | BCS1 homolog, ubiquinol-cytochrome c reductase complex chaperone | 301514 |
| Casp3 | Caspase 3 | 25402 |
| Casp8 | Caspase 8 | 64044 |
| Col1a1 | Collagen alpha-1(I) chain | 29393 |
| Col3a1 | Collagen alpha-1(III) chain | 84032 |
| Cox4i1 | Cytochrome c oxidase subunit 4i1 | 29445 |
| MMP2 | Matrix metalloproteinase 2 | 81686 |
| MMP9 | Matrix metalloproteinase 9 | 84687 |
| MT-ATP6 | ATP synthase 6, mitochondrial | 26197 |
| Mybl2 | MYB proto-oncogene like 2 | 296344 |
| Ndufa1 | NADH:ubiquinone oxidoreductase subunit A1 | 363441 |
| Pcna | Proliferating cell nuclear antigen | 25737 |
| Sdha | Succinate dehydrogenase complex flavoprotein subunit A | 157074 |
| Tfgbr3 | Transforming growth factor beta receptor 3 | 29610 |
| TIMP1 | Tissue inhibitor of matrix metalloproteinase 1 | 116510 |

### Mitochondrial DNA Content

mtDNA content was quantified by ratio of a target mitochondrial gene, NADH dehydrogenase subunit 1 (ND1), to nuclear reference gene, protein argonaute-1 (EIF2C1)^62, 63^. Total DNA was isolated using the Quick-DNA Microprep kit (Zymo Research, Irvine, CA). DNA concentration was determined using a NanoDrop 2000 (Thermo Scientific) and serial diluted to 1ng/μl. A droplet digital PCR (ddPCR) method was used to complete quantification of absolute copy number of both ND1 and EIF2C1. Reactions were prepared in duplex format, in a final volume of 20μl with 2x ddPCR Supermix for Probes (No dUTP) (BioRad, Hercules, CA), 20x reference probe ND1 labeled with FAM (BioRad), 20x reference probe EIF2C1 labeled with HEX (BioRad), 5ng of DNA, 1μl of HindIII enzyme (FastDigest, Thermo Scientific), and nuclease-free water. The assembled reaction was incubated at room temperature for 20 minutes to permit digestion with the HindIII restriction enzyme. After digestion, droplets were generated from prepared PCR reactions using Droplet Generation Oil for Probes (BioRad) on a QX200 Droplet Generator (BioRad), transferred to a deep-well 96-well plate, and amplified (95°C-10 minutes, 1 cycle; 94°C-30 seconds and 60°C-90 seconds, 40 cycles; 98°C-10 minutes, 1 cycle) on a C1000 thermal cycler (BioRad). At completion of end-point PCR, absolute quantification of PCR products was completed on a QX200 Droplet Reader (BioRad) with QuantaSoft Software Version 1.7 (BioRad) and reported as a ratio of positive ND1/EIF2C1 counts per 20μl reaction.

### Protein Expression

Cultured cells were lysed in RIPA buffer containing 50mM Tris-HCl, 150mM NaCl, 2mM EDTA, 2mM EGTA, 0.5% sodium deoxycholate, 1% Triton X-100, 0.1% SDS, 50mM NaF, 0.2mM Na_3_VO_4,_ and 0.2% protease inhibitor cocktail (Sigma). Samples were prepared as described previously^64^. Equal amounts of protein (15μg) were separated in duplicate on an Any-kD TGX polyacrylamide gel (BioRad). Protein was transferred to a PVDF membrane (TransBlot Turbo, BioRad) and blocked for 24 hours at 4°C in 5% blotting-grade blocker (BioRad). Blots were incubated in primary OXPHOS antibody (1:1000, 110413, Abcam, Cambridge, MA) and then HRP-conjugated goat anti-mouse antibody (1:5000, 31430, Invitrogen), each for 2 hours at room temperature. Bands of interest were compared against positive control (Rat Heart Mitochondria, Abcam) and all targets were probed on the same membrane, but required different exposure times using signal accumulation mode (ChemiDoc, BioRad). Volume intensity analysis of matched exposure duration for each target was completed using Image Lab Version 6.0.1 (BioRad). Total protein volume intensity obtained by UV activated Stain-Free imaging was used for data normalization.

### Superoxide Production

Superoxide (O_2_^-^) production was determined by dihydroethidium (DHE) derived fluorescence. Cultured cells were detached, pelleted, and resuspended in Hank’s Balanced Salt Solution. An aliquot was saved for determination of cell concentration. DHE (Thermo Scientific) was added at final concentration of 10μM and samples were incubated with shaking at 37°C for 30 minutes^65–67^. Fluorescent spectrum was recorded at 530nm excitation/590nm emission (Molecular Devices) and fluorescence units were normalized to cell concentration.

### Statistical Analysis

Statistical analyses on the four main effect contrasts of interest (50μg/ml AGE vs. 0μg/ml AGE, 100μg/ml AGE vs. 0μg/ml AGE, 200μg/ml AGE vs. 0μg/ml AGE, and HG vs. LG) for each outcome variable proceeded via a mixed effects regression model with consideration of technical data replicates and random effects to account for the tendon fibroblast donor rats. Regression models were fit to the data using the `lmer` function in `R`. The Kenward-Roger F-test was performed to determine whether the main effects regression model or the “full” model containing both main effects and two-factor interactions between the AGE and glucose contrasts should be fitted. Validity of the assumptions for the models were assessed via standard regression diagnostics. Specifically, the standardized residuals were examined to assess whether they were approximately Normally distributed, centered at zero, and did not exhibit any patterns with respect to the predictor variables. If assumptions appeared invalid for the original scale, the logarithmic and square root transformations were considered. After identifying a model that provided a good fit to the outcome data, hypothesis tests were performed for the contrasts using Satterthwaite’s method, and confidence intervals were created using the bootstrap. Hypothesis tests for the contrasts use a Bonferroni-adjusted significance level of α=0.05/4=0.0125 (Figures 1-5 and 7) or α=0.05/3=0.017 (Figures 6 and 8) separately across the different outcome variables to account for the multiple comparisons. Certain specified, direct comparisons between two specific groups were performed via the paired t-test in `R` with α=0.05 to test conditional main effects, where HG was conditioned for testing the AGE contrast of interest. Summary of applied contrasts and exact p-values for significant findings are provided in Table 2. Figures were generated in Prism 7.0 (GraphPad, La Jolla, CA) and are represented as mean ± standard error.

**Table 2:** Summary of p-values for Statistical Findings

| Figure | AGE-0µg/ml vs.<br>AGE-50µg/ml | AGE-0µg/ml<br>vs.<br>AGE-100µg/ml | AGE-0µg/ml<br>vs.<br>AGE-200µg/ml | Normal vs.<br>High Glucose | Interaction |
| --- | --- | --- | --- | --- | --- |
| 1B | $4 \times 10^{-5}$ | $3 \times 10^{-14}$ | $< 2 \times 10^{-16}$ | NS | N/A |
| 1C | $1 \times 10^{-8}$ | $5 \times 10^{-15}$ | $< 2 \times 10^{-16}$ | NS | N/A |
| 1D | $1.5 \times 10^{-10}$ | $< 2 \times 10^{-16}$ | $< 2 \times 10^{-16}$ | NS | N/A |
| 1E | $4 \times 10^{-9}$ | $2 \times 10^{-15}$ | $< 2 \times 10^{-16}$ | $7 \times 10^{-3}$ | N/A |
| 1F | $5 \times 10^{-9}$ | $1.5 \times 10^{-14}$ | $< 2 \times 10^{-16}$ | $8 \times 10^{-4}$ | N/A |
| 2A | NS | $3 \times 10^{-7}$ | $2 \times 10^{-14}$ | $1 \times 10^{-3}$ | N/A |
| 2B | NS | $9 \times 10^{-6}$ | $5 \times 10^{-10}$ | NS | N/A |
| 2C | NS | NS | NS | NS | N/A |
| 2D | NS | NS | NS | $2 \times 10^{-3}$ | N/A |
| 2E | NS | NS | $1 \times 10^{-14}$ | $1 \times 10^{-7}$ | N/A |
| 2F | NS | NS | $3 \times 10^{-4}$ | NS | N/A |
| 3A | NS | NS | $5 \times 10^{-11}$ | NS | N/A |
| 3B | $5 \times 10^{-4}$ | $1.4 \times 10^{-5}$ | NS | NS | N/A |
| 3C | NS | $3 \times 10^{-4}$ | NS | NS | N/A |
| 3D | NS | NS | $7 \times 10^{-3}$ | NS | N/A |
| 3E | NS | NS | NS | NS | N/A |
| 4A | $6 \times 10^{-7}$ | $1 \times 10^{-12}$ | $2 \times 10^{-15}$ | NS | N/A |
| 4B | NS | NS | NS | NS | N/A |
| 5A | $6 \times 10^{-6}$ | $8 \times 10^{-7}$ | $7 \times 10^{-5}$ | NS | N/A |
| 5B | NS | NS | $5 \times 10^{-3}$ | NS | N/A |
| 5C | $2 \times 10^{-5}$ | $1 \times 10^{-11}$ | $< 2 \times 10^{-16}$ | $1 \times 10^{-2}$ | N/A |
| 5D | NS | NS | NS | NS | N/A |
| 5E | NS | NS | NS | NS | N/A |
| 6A | N/A | N/A | NS | NS | $1.2 \times 10^{-2}$ |
| 6B | N/A | N/A | NS | NS | NS |
| 6C | N/A | N/A | $1.3 \times 10^{-2}$ | NS | NS |
| 6D | N/A | N/A | NS | NS | $2.8 \times 10^{-3}$ |
| 6E | N/A | N/A | NS | NS | NS |
| 7A | NS | $9 \times 10^{-7}$ | $< 2 \times 10^{-16}$ | NS | N/A |
| 7B | NS | NS | NS | NS | N/A |
| 7C | NS | NS | $7 \times 10^{-10}$ | NS | N/A |
| 7D | NS | NS | NS | $1.1 \times 10^{-2}$ | N/A |
| 7E | NS | NS | NS | NS | N/A |
| 7F | NS | NS | $1 \times 10^{-4}$ | NS | N/A |
| 8 | N/A | N/A | $8.6 \times 10^{-3}$ | NS | NS |

NS = Not Significant
N/A = Not Tested

## Acknowledgements

This work was supported by NIH pre-doctoral fellowship F31-AR073647 (SHP) and Purdue University Research Initiative Funds (CCC). The authors would like to acknowledge Zachary Hettinger, MS, Christopher Kargl, MS, and Sukhmani Kaur for technical laboratory assistance.

## Author Contributions

S.H.P and C.C.C conceived and designed the study. S.H.P, F.Y, S.K.S, R.F, A.S, and C.C.C performed data collection and analysis. S.H.P, F.Y, S.K.S, R.F, J.R.C, J.H.S, S.K, A.S, and C.C.C interpreted data and provided expert advice. S.H.P and C.C.C drafted the manuscript. All authors edited, revised, read, and approved the final submitted manuscript.

## Competing Interests

The authors declare no competing interests.

## Data Availability

C.C.C has access to all data and data is available upon request.

